# Sumoylation is largely dispensable for normal growth but facilitates heat tolerance in yeast

**DOI:** 10.1101/761759

**Authors:** Marjan Moallem, Akhi Akhter, Giovanni L. Burke, John Babu, Benjamin G. Bergey, J. Bryan McNeil, Mohammad S. Baig, Emanuel Rosonina

**Author notes:** Address correspondence to Emanuel Rosonina.

## Abstract

Numerous proteins are sumoylated in normally growing yeast and SUMO conjugation levels rise upon exposure to several stress conditions. We observe high levels of sumoylation also during early exponential growth and when nutrient-rich medium is used. However, we find that reduced sumoylation (~75% less than normal) is remarkably well-tolerated, with no apparent growth defects under non-stress conditions or under osmotic, oxidative, or ethanol stresses. In contrast, strains with reduced activity of Ubc9, the sole SUMO conjugase, are temperature-sensitive, implicating sumoylation in the heat stress response, specifically. Aligned with this, a mild heat shock triggers increased sumoylation which requires functional levels of Ubc9, but likely also depends on decreased desumoylation, since heat shock reduces protein levels of Ulp1, the major SUMO protease. Furthermore, we find that a *ubc9* mutant strain with only ~5% of normal sumoylation levels shows a modest growth defect, has abnormal genomic distribution of RNA polymerase II (RNAPII), and displays a greatly expanded redistribution of RNAPII after heat shock. Together, our data implies that SUMO conjugations are largely dispensable under normal conditions, but a threshold level of Ubc9 activity is needed to maintain transcriptional control and to modulate the redistribution of RNAPII and promote survival when temperatures rise.

## INTRODUCTION

Nearly one tenth of all human and budding yeast proteins have been identified as targets of SUMO post-translational modification in non-stress growth conditions, a large fraction of which are involved in gene expression and chromatin maintenance or regulation (1–6). The effects of protein sumoylation are largely target-specific, and include altered localization, activity, stability, and association with other proteins and chromatin (5–9). The context-specific nature of the consequences of sumoylation is exemplified by its apparently contrasting roles in regulating transcription (9, 10). Whereas tethering SUMO or Ubc9, the SUMO conjugating enzyme, to the promoters of reporter genes dramatically reduced their expression, genome-wide chromatin immunoprecipitation (ChIP) studies showed that sumoylated proteins are found predominantly at promoters of transcriptionally active genes (7, 11–16). Furthermore, hundreds of sequence-specific transcription factors (SSTFs) are known SUMO conjugates, and while their sumoylation is frequently associated with reduced expression of target genes, efficient gene activation by some SSTFs has been shown to depend on their sumoylation (8). Sumoylation, therefore, does not appear to have general transcription-promoting or transcription-repressing roles, but instead it likely controls expression of numerous genes in different ways (8, 9, 17, 18).

SUMO conjugation involves the covalent attachment of a SUMO peptide to the side chain of specific Lys residues, through isopeptide bonds, that often lie within the SUMO consensus motif (1, 7). It occurs through a cascade of enzymatic activities that is analogous to ubiquitination: activation by an E1 class enzyme and target conjugation by E2 enzymes, which can be facilitated by E3 SUMO ligases that enhance sumoylation and target Lys specificity (7, 19, 20). Although many proteins are sumoylated, typically only a small fraction (<5%) of polypeptides of each target protein is modified at any one time, which is partly the result of the constitutive activity of SUMO proteases, including the SENP family of isopeptidases in human cells (21). In yeast, there are two SUMO proteases, Ulp1, which is essential for viability, and Ulp2, which prevents accumulation of polysumoylate chains that form when conjugated SUMO peptides themselves become sumoylated (22–24). In sharp contrast with ubiquitination, the number of SUMO conjugation and deconjugation enzymes is small, with both yeast and mammals harboring only a single E2 conjugase, Ubc9. As such, eukaryotic cells can simultaneously control the sumoylation level of hundreds of substrate proteins by regulating the activity of just one SUMO conjugating or deconjugating enzyme at a time. For example, exposure of yeast to ethanol triggers the nucleolar sequestration of Ulp1, which results in a global increase in protein sumoylation within minutes (25).

Such rapid increases in global levels of SUMO conjugation occur in response to a variety of stress conditions, and are collectively referred to as the SUMO stress response (SSR; (17, 18, 26–28)). Exposure of mammalian cells to heat, oxidative, osmotic, or ethanol stress results in a spike in sumoylation levels caused by increased conjugation of numerous proteins with SUMO isoforms SUMO2 and/or SUMO3, specifically (29–31). As with sequestration of Ulp1 in yeast upon exposure to ethanol, the SSR that occurs with heat shock in human cells is at least partly caused by inactivation of multiple SENPs, suggesting that constitutive desumoylation is required for maintaining steady state levels of sumoylation (32). In addition to ethanol exposure, yeast also respond with dramatically elevated SUMO conjugation levels in response to oxidative and osmotic conditions (26, 33). Inhibitors of transcription virtually eliminate the SSR in yeast, implying that elevated sumoylation is largely a consequence of stress-induced transcriptional reprogramming and is not necessarily linked to the stress itself (26). However, SUMO2/3 isoforms are required for human cells to survive heat shock, and elevated levels of sumoylation appear to have a protective effect on neuronal tissue during exposure to reduced oxygen and glucose (7, 31, 34–36). This suggests that elevated sumoylation itself can serve a role in the adaptation of cells to unfavourable conditions, but the physiological effects of modulated cellular sumoylation levels are largely unknown.

To explore this, here we examined how reduced sumoylation affects cell fitness and transcription patterns in yeast. Our data show that most SUMO modifications are not needed for normal growth under non-stress and some stress conditions, but that active SUMO conjugation by Ubc9 is critical when temperatures rise. Moreover, we find that a mild heat shock triggers elevated sumoylation levels at least partly by reducing the abundance of Ulp1. Mutant yeast with dramatically reduced sumoylation levels display widespread changes to RNA polymerase II (RNAPII) occupancy across protein-coding genes in normal growth, and the heat-shock triggered redistribution of RNAPII is greatly exacerbated in this strain. Taken together, our results suggest that, whereas some level of constitutive sumoylation is required to maintain normal transcription patterns, sumoylation and desumoylation systems play largely pre-emptive roles in preparing yeast for potential exposure to elevated temperature.

## RESULTS

### Cellular sumoylation levels are modulated under different conditions

Proteomics studies have identified nearly 600 SUMO conjugates in budding yeast (2–4), and many of these can be detected by SUMO immunoblot analysis. For example, cell lysates from normally growing, unstressed yeast of the common lab strain W303a were analyzed by immunoblot with the y-84 SUMO antibody, which showed multiple bands as well as unconjugated (“free”) SUMO, as shown in Fig. 1A. The detection level of SUMO conjugation was reduced when the SUMO protease inhibitor *N*-ethylmaleimide (NEM) was excluded from lysate preparations, and sumoylation patterns were shifted upwards on the immunoblot when a strain expressing SUMO with an 8xHIS N-terminal epitope tag was used, confirming that the antibody recognizes SUMO and SUMO conjugates, specifically. Intriguingly, the antibody also strongly detected a sumoylated species migrating at ~30 kDa, which most likely corresponds to Ubc9 modified by SUMO (“Ubc9-S” in diagrams) since (i) it co-migrates with Ubc9-SUMO as detected in a Ubc9 immunoblot (Fig. 1A, right), (ii) its migration in the 8HIS-Smt3 strain is retarded to a similar degree in the SUMO and Ubc9 blots, and (iii) fusion of polypeptide sequences to Ubc9 results in a corresponding upward shift in migration of this species (see below). Specifically, this species likely derives from Ubc9 modified with SUMO through a Lys-linked isopeptide bond, not through a thioester bond to Cys, since it is resistant to highly reducing conditions (Fig. S1A). Altogether, this analysis confirms that we can detect high levels of protein sumoylation in normally growing yeast in addition to unconjugated SUMO and sumoylated Ubc9.

**Figure 1.**
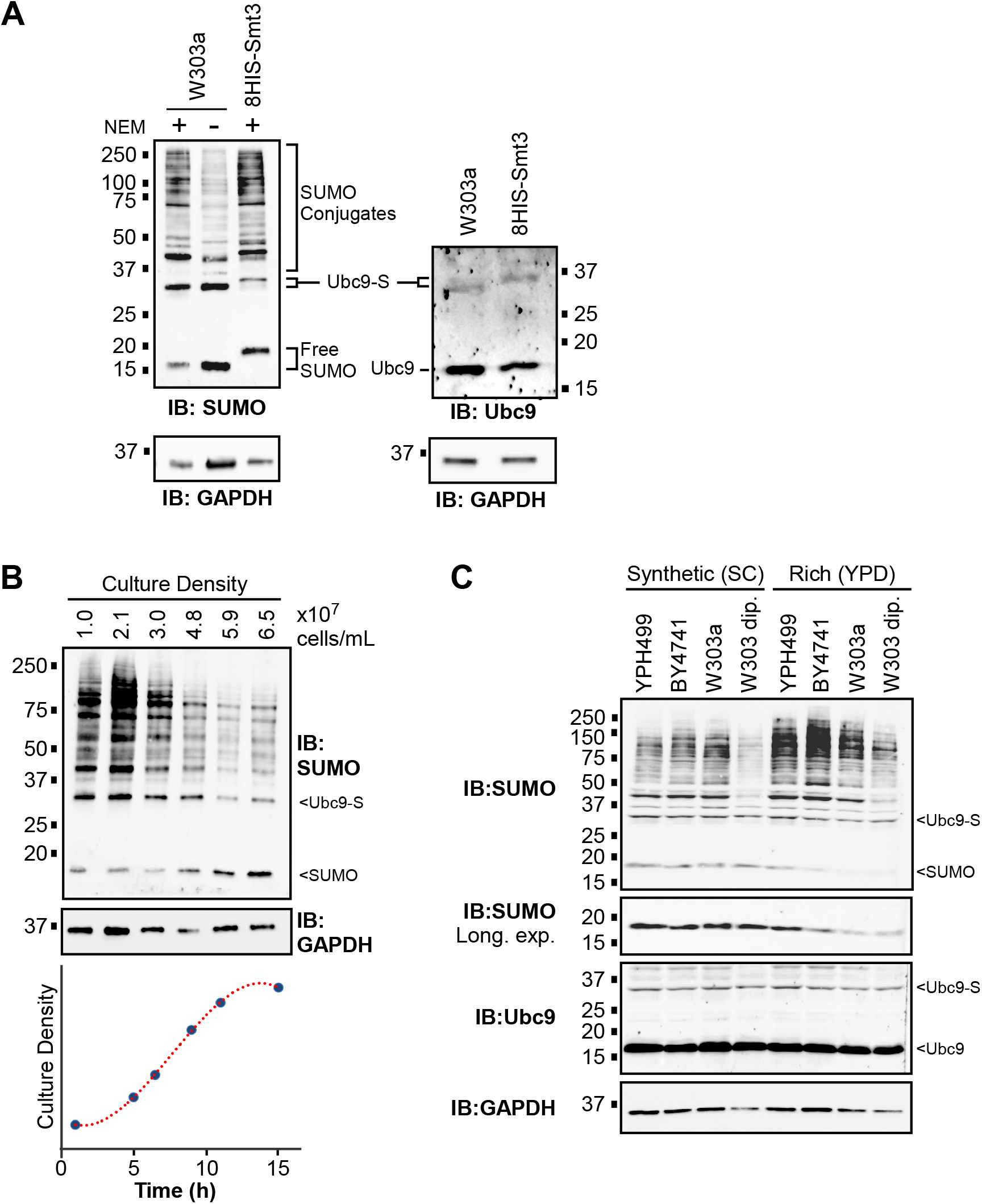
Many proteins are dynamically sumoylated in non-stressed yeast. **(A)** W303a and 8HIS-Smt3 yeast strains were grown in SC medium at the standard temperature (30°C) and cell lysates were prepared under non-denaturing conditions, in the presence of NEM, except where indicated. Lysates were analyzed by SUMO, GAPDH, and Ubc9 immunoblots under conditions that facilitate detection of smaller proteins (see Experimental Procedures). The positions of SUMO conjugates, Ubc9-SUMO (“Ubc9-S”), unconjugated SUMO (“Free SUMO”), and unmodified Ubc9 are indicated. **(B)** Aliquots of a W303a culture grown in SC medium at 30°C were collected at six intervals over 15 hours and protein extracts were prepared and analyzed by SUMO and GAPDH immunoblots. The culture density (cells/mL) at each time point is indicated at top and depicted in the curve below. **(C)** The indicated common lab yeast strains, including haploid and diploid W303 (“W303a” and “W303 dip.,” respectively), were grown in either SC or YPD medium to exponential phase, then protein extracts were generated and analyzed by SUMO, Ubc9, and GAPDH immunoblots.

Next, we tested whether factors other than the previously reported stress conditions can affect SUMO conjugation levels. The W303a strain was grown in liquid culture and protein extracts were generated from aliquots taken from the culture at various time points and examined by SUMO immunoblot. Remarkably, the level of sumoylation dropped dramatically as the culture density increased, suggesting that depletion of nutrients, or physiological changes that accompany the approach to culture saturation (e.g., diauxic shift), results in reduced SUMO conjugation levels (Fig. 1B). Finally, we compared sumoylation levels in different common lab strains of yeast grown in either synthetic (SC) or nutrient-rich (YPD) media. Minor differences were seen between the YPH499, BY4741, and W303a haploid yeast strains, but significantly less sumoylation was detected in a diploid yeast strain (W303 diploid), and approximately two times more sumoylation was detected when strains were grown in YPD versus SC medium (Fig. 1C). Together, our results indicate that yeast cells modulate cellular sumoylation levels in response to culture density, nutrient availability (i.e., medium “richness”), ploidy, and to a lesser extent, strain genetic background.

### Yeast tolerate dramatically reduced sumoylation levels with no apparent growth defect

To explore the physiological consequences of reduced global sumoylation, we applied the Anchor Away methodology to Ubc9 (37). By this system, Ubc9 is fused with the FRB domain of human mTOR (“Ubc9-AA”) in a yeast strain that also expresses the ribosomal protein Rpl13A fused to human FKBP12, which binds FRB in the presence of rapamycin. Exposing the Ubc9-AA strain to rapamycin causes nuclear Ubc9-AA to bind Rpl13A-FKB12, which is then rapidly translocated to the cytoplasm, effectively depleting the nucleus of Ubc9 and de novo sumoylation. As most sumoylated proteins are nuclear and associated with chromatin (16), this is expected to cause a significant reduction in overall cellular sumoylation levels. The Anchor Away parent strain, Parent-AA, harbours a mutation in the *TOR1* gene such that exposure to rapamycin has no effect on growth or sumoylation levels (see Figs. 2A, C, D). We compared sumoylation levels in the Parent-AA and Ubc9-AA strains, which are isogenic except for the C-terminal FRB fusion on Ubc9 in the latter. Notably, even in the absence of rapamycin, about 75% less sumoylation was detected in the Ubc9-AA strain, indicating that the FRB fusion itself hampers Ubc9 activity (Fig. 2A). This might reflect a lower abundance of Ubc9-AA compared to normal Ubc9, as detected in a Ubc9 immunoblot (Fig. 2B), but we cannot rule out that the fusion inhibits binding of the Ubc9 antibody, which recognizes the C-terminus of Ubc9. Although sumoylation levels are reduced, the Ubc9-AA strain responds normally to ethanol, oxidative, and osmotic stress with increased SUMO conjugation (Fig. S1B; (26, 33)). Addition of rapamycin for 30 min during growth caused a further dramatic reduction in overall sumoylation levels (to ~3% of parental levels) in the Ubc9-AA strain, which is consistent with the exclusion of Ubc9-AA from the nucleus in these conditions (Fig. 2A). The Ubc9-AA system, therefore, allows us to examine the effects of constitutively low SUMO conjugation levels which can be further reduced by exposure to rapamycin.

**Figure 2.**
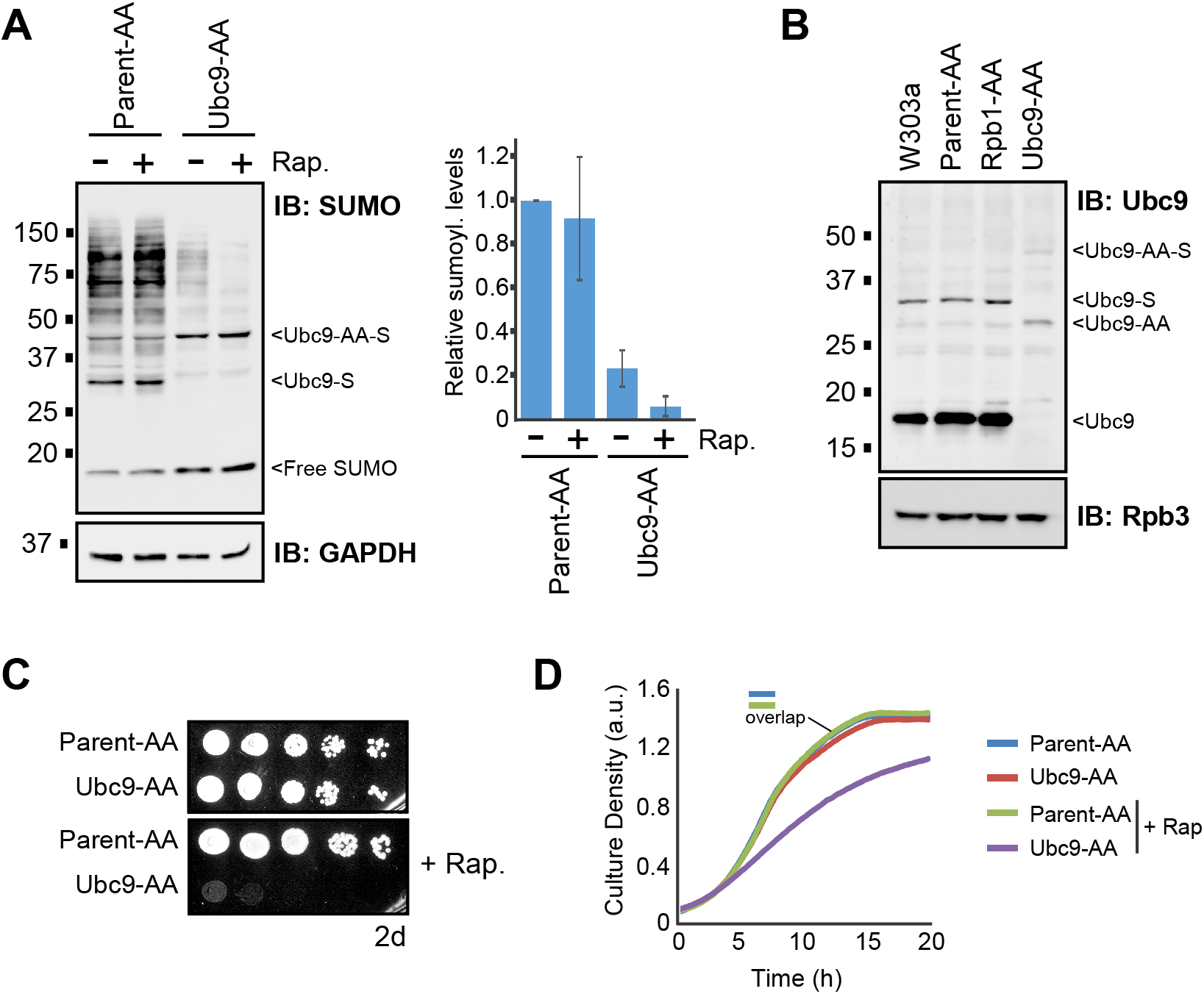
Yeast display very high tolerance for reduced sumoylation levels in non-stress growth conditions. **(A)** Cultures of the Ubc9 Anchor Away strain, Ubc9-AA, and its parent, Parent-AA, were left untreated or treated with rapamycin (“+Rap.”) for 30 min, then extracts were prepared and analyzed by SUMO and GAPDH immunoblots. SUMO conjugation levels were quantified by densitometry and the average and standard deviations (error bars) from three experiments are plotted at right. **(B)** Lysates were prepared from cultures of W303a, Parent-AA, Ubc9-AA, or from an unrelated control Anchor Away strain, Rpb1-AA, and analyzed by immunoblots with antibodies that recognize Ubc9 or a subunit of RNA Polymerase II, Rpb3, which serves as a loading control. **(C)** A spot assay was performed with the Ubc9-AA and Parent-AA strains, on either SC medium, or SC supplemented with rapamycin. Plates were imaged after two days of growth. **(D)** Cultures of the Ubc9-AA and Parent-AA strains were prepared in SC medium, with or without rapamycin, at an absorbance (595 nm) of ~0.2, then grown for 20 hours while absorbance (culture density) measurements were made every 15 min. The two Parent-AA curves are virtually indistinguishable and are therefore marked with “overlap.” Triplicate cultures were grown for each sample, and the average values were plotted to create the growth curves shown. Values for average absorbance measurements and standard deviations for each triplicate set are listed in Table S8.

We then compared growth rates of the Parent-AA and Ubc9-AA strains in the presence or absence of rapamycin. Strikingly, despite the ~75% reduction in sumoylation levels in Ubc9-AA, the strains grew equally well in untreated SC medium, indicating that normal levels of SUMO conjugation are not needed for growth in standard conditions on solid medium (Fig. 2C) or during exponential growth in liquid culture (Fig. 2D). Further reducing sumoylation levels by the addition of rapamycin, however, was lethal to the Ubc9-AA strain, as indicated by spot assay (Fig. 2C). This effect is supported by the liquid growth assay that showed significantly slowed escape from lag phase and attenuated exponential growth for Ubc9-AA in the presence of rapamycin, which had no effect on the Parent-AA strain (Fig. 2D). As also shown in experiments described below, we note that small differences in our growth curves, specifically at the level of cell saturation, generally relate to much more dramatic growth defects in spot assays. In any case, our data indicate that yeast growth is surprisingly unaffected by a dramatic reduction in cellular sumoylation levels, although levels of ~3% are lethal.

### Yeast with constitutively low Ubc9 activity show high temperature sensitivity

To determine whether the high tolerance for reduced sumoylation can be observed in other contexts, we examined the *ubc9-6* and *ubc9-1* mutant strains and their respective isogenic parental strains (16, 38). The *ubc9-6* and *ubc9-1* strains harbor a point mutation that partly inactivates Ubc9, and like Ubc9-AA, they show constitutively low levels of sumoylation, with *ubc9-6* showing the most dramatic reduction (~5%) compared to its parent, W303a (Fig. 3A). Consistent with this, spot assays and liquid growth curves showed that only *ubc9-6* shows a growth defect, albeit modest, in standard growth conditions, which supports the idea that only a very low threshold level of sumoylation is needed for normal growth (Fig. 3B, C).

**Figure 3.**
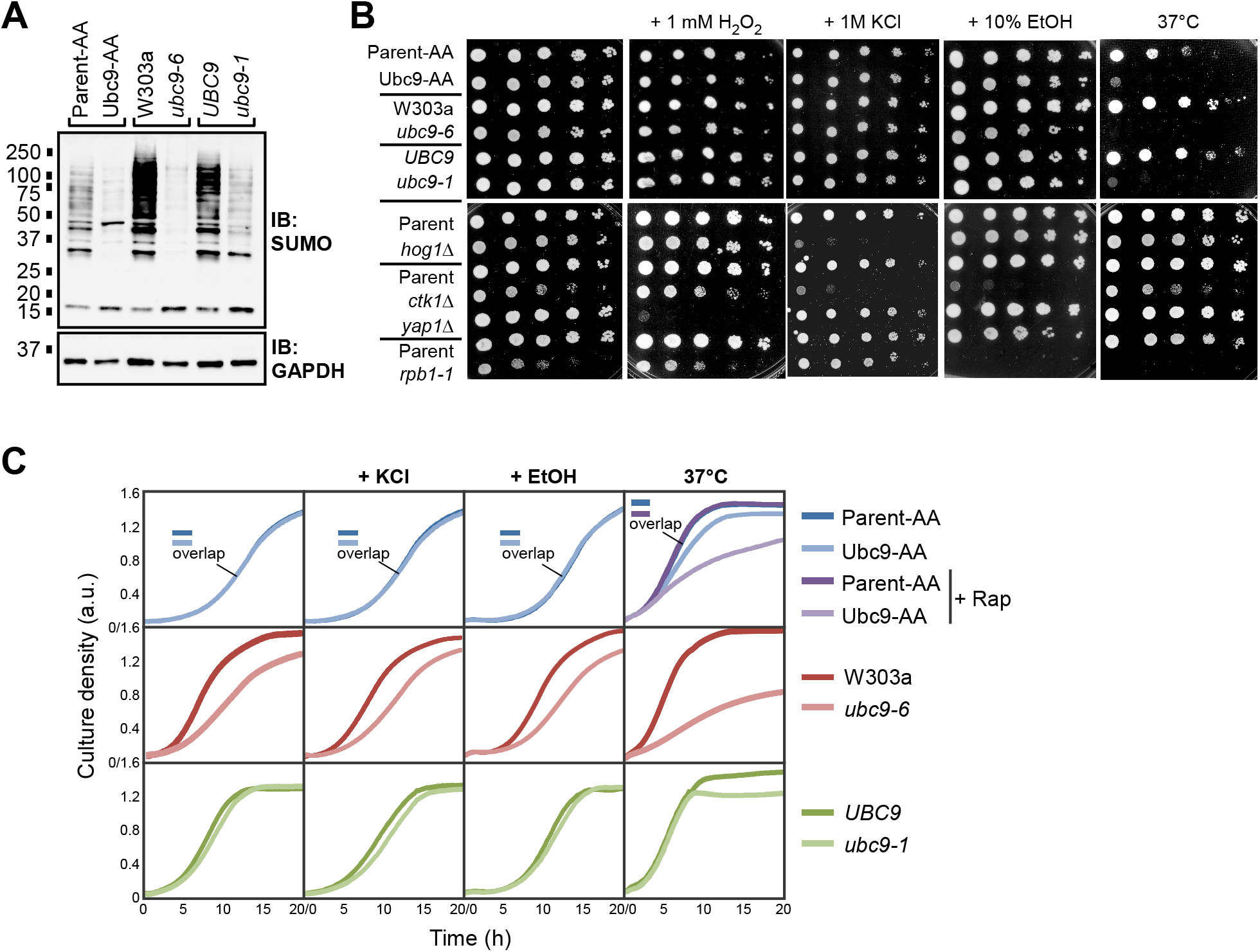
Yeast strains with constitutively low levels of Ubc9 activity cannot survive elevated temperatures. **(A)** Protein extracts were prepared from cultures of the indicated strains grown in untreated SC medium and analyzed by SUMO and GAPDH immunoblots. **(B)** Spot assays were performed with the indicated strains, onto either SC medium, or SC supplemented with the indicated stressors (see Experimental Procedures) and incubated at the standard growth temperature of 30°C, or at the elevated temperature of 37°C. **(C)** Growth curves were generated, as in Fig. 2D, for the indicated strains and conditions. Curves that overlap significantly and are indistinguishable are marked with “overlap.” Individual graphs include two curves except for the Parent-AA and Ubc9-AA set at 37°C, which includes untreated and rapamycin-treated samples for both strains.

Why do cells maintain levels of sumoylation that are significantly higher than needed for normal growth? Considering that sumoylation levels increase in response to oxidative, osmotic, ethanol, and temperature stresses (26–28, 33), we wished to determine whether normal sumoylation levels might play a role in preparing cells for possible exposure to these conditions. Surprisingly, the Ubc9-AA, *ubc9-6*, and *ubc9-1* strains grew as well on medium containing H2O2, KCl, or ethanol as they did on standard medium, relative to their parental strains (Fig. 3B). This was also observed in liquid growth assays in media containing KCl or ethanol (Fig. 3C), and it indicates that normal levels of sumoylation are not required for survival, or even optimal growth, through these stress conditions. In sharp contrast, all three mutant strains showed a strong growth defect when grown at the elevated temperature of 37°C on solid medium (Fig. 3B), and significantly impaired growth in the liquid culture assay (Fig. 3C), with *ubc9-6* showing the most severe defect (Fig. 3C). The *ubc9-6* and *ubc9-1* strains were previously characterized as temperature-sensitive (16, 38), and it was suggested that this sensitivity is due to instability of mutant Ubc9 at elevated temperature (see below). However, also considering the temperature-sensitivity of the Ubc9-AA strain, one intriguing possibility is that these strains are unable to grow well at 37°C because sumoylation is required to survive when temperatures increase.

### Yeast with induced loss of Ubc9 also grow normally except at high temperatures

To examine the effects of reduced cellular sumoylation levels using an additional approach, we employed the Ubc9-TO (“Tet-Off”) strain, in which the *UBC9* gene is under the control of a tetracycline repressible promoter (39). The strain was grown for several hours in medium containing varying concentrations of the tetracycline analog doxycycline, then lysates were prepared and analyzed by SUMO and Ubc9 immunoblots. Even at a low concentration of doxycycline (0.5 μg/mL), both Ubc9 and SUMO conjugation showed a dramatic reduction to less than 5% of their normal levels (Fig. 4A). We then examined how well the Ubc9-TO strain tolerates low sumoylation levels, and whether reduced sumoylation also confers temperature sensitivity in this strain. Despite the dramatic reduction in sumoylation levels, treatment with doxycycline had no effect on exponential growth of the Ubc9-TO strain at 30°C, which is again consistent with our finding that yeast cells show high tolerance for low levels of sumoylation when grown at optimal temperatures (Fig. 4B). When the analysis was performed at elevated temperatures, however, Ubc9-TO cells treated with doxycycline displayed a growth defect which was more pronounced at 39.5°C than at 37°C, whereas the drug had no effect on the isogenic Parent-TO strain at any temperature (Fig. 4B). Using this different, inducible system, these observations support our finding that many SUMO conjugation events are dispensable under optimal growth conditions but are needed to prepare cells for elevated temperatures.

**Figure 4.**
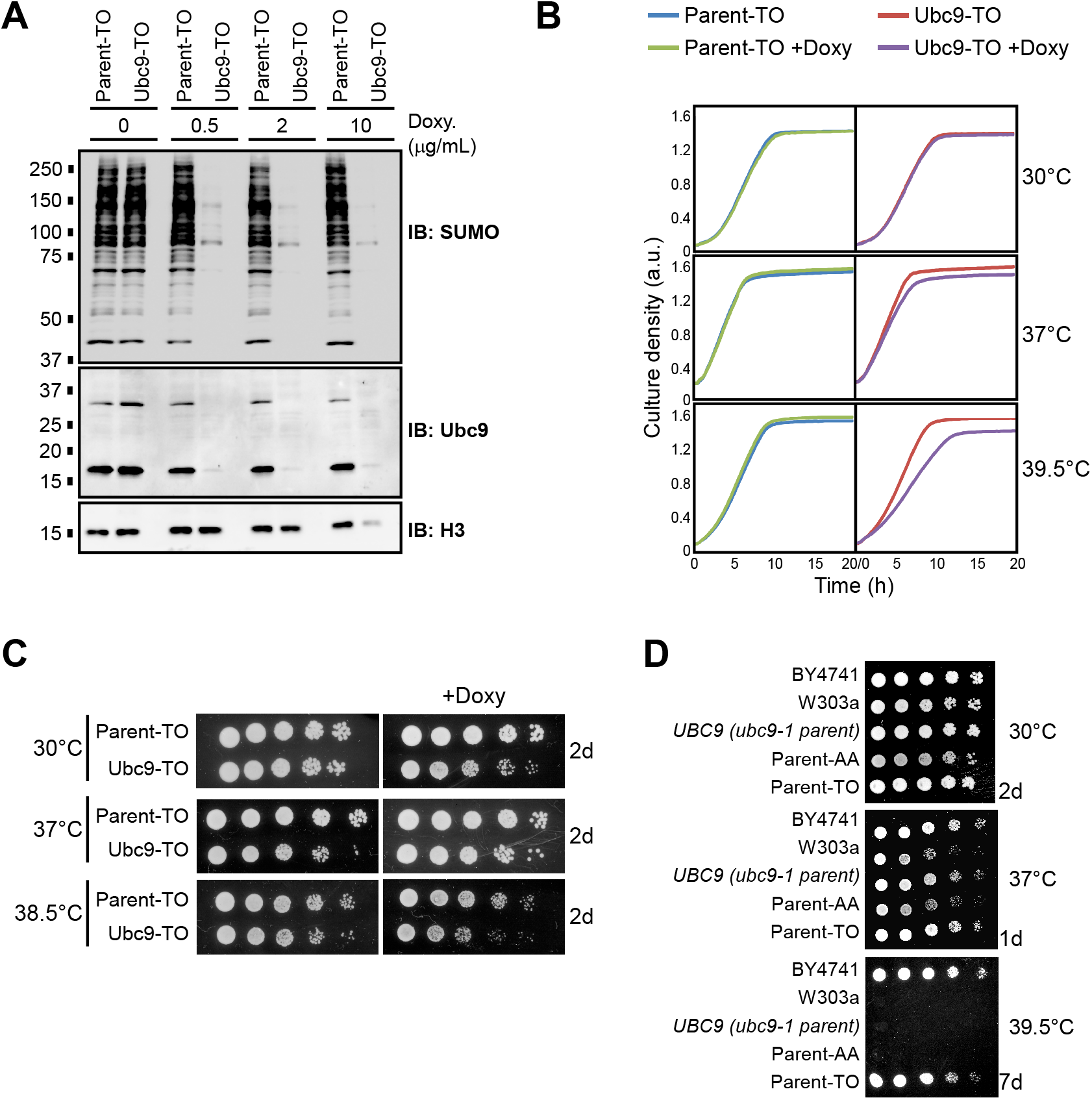
Induced reduction of Ubc9 levels also sensitizes yeast to high temperatures. **(A)** Cultures of a strain expressing the *UBC9* gene from a tetracycline-repressible promoter, Ubc9-TO, and its isogenic parent strain, Parent-TO, were exposed to the indicated concentrations of the tetracycline analog doxycycline for several hours, then lysates were prepared and examined by SUMO, Ubc9, and H3 immunoblots. **(B)** Growth curves were generated, as in Fig. 2D, for the Ubc9-TO and Parent-TO strains grown in the absence or presence of doxycycline at the indicated temperatures. **(C)** Spot assays were performed with the Ubc9-TO and Parent-TO strains, onto either SC medium, or SC supplemented with doxycycline. Plates were incubated at the indicated temperatures and were imaged after two days of growth. As none of the strains grow in the presence of doxycycline at 39.5°C on solid medium, a temperature of 38.5°C was used as the highest temperature for this experiment. **(D)** A spot assay was performed comparing growth of the indicated background and parental yeast strains at 30°C, 37°C, or 39.5°C, for the indicated number of days.

We also examined how reduced sumoylation affects growth of the Ubc9-TO strain on solid medium using spot assays. Parent-TO and Ubc9-TO strains were grown in culture for several hours, either untreated or in the presence of doxycycline, then spotted onto the same medium in solid form. Compared to the Parent-TO strain, a modest growth defect was observed in the Ubc9-TO strain when grown in the presence of doxycycline at 30°C, suggesting that the dramatic reduction in sumoylation has some impact on growth on solid medium (Fig. 4C). The growth defect was more apparent when samples were incubated at 38.5°C, which was the highest temperature at which either strain could grow on doxycycline-containing solid medium (Fig. 4C). As the Ubc9-TO and Parent-TO strains derive from a different background yeast strain (BY4741) than the Ubc9-AA and Parent-AA strains (W303a), we explored whether yeast strains of different origins show differential sensitivity to high temperatures. Indeed, as shown in Fig. 4D, BY4741, and its derived strain Parent-TO, show strikingly higher thermotolerance than W303a-derived strains, including the Parent-AA strain and the parental strain for *ubc9-1, UBC9*. Although this inherent high tolerance for elevated temperatures may obscure temperature sensitivity to reduced sumoylation levels in the Ubc9-TO strain when spot assays are performed, the liquid growth assays shown in Fig. 4B support our finding that yeast cells with reduced SUMO conjugation levels show growth defects at high temperatures.

### Mild heat shock elevates global sumoylation levels and reduces Ulp1 protein abundance

Previous studies showed that sumoylation levels increase in response to heat shock in human cells and to a high-temperature heat shock (42°C) in yeast (28, 31, 32). To examine this in further detail, wild-type W303a yeast cultures were exposed to a mild heat shock (37°C) for increasing durations, lysates were prepared, then examined by SUMO immunoblot. Indeed, high molecular-weight SUMO conjugation levels increased through the time-course and peaked at 30 min with about 50% more sumoylation (Fig. 5A). In contrast, no increase in sumoylation was observed in the Ubc9-AA, *ubc9-6*, or *ubc9-1* strains after a 30-min heat shock at 37°C (Fig. 5B). Potentially explaining this, Ubc9 protein levels decreased in the *ubc9-6* and *ubc9-1* strains during the heat shock, and Ubc9 levels in the Ubc9-AA strain are constitutively very low (Figs. 5B and 2B). Although the further reduction in Ubc9 levels in the *ubc9-6* and *ubc9-1* strains during heat shock may contribute to the temperature-sensitivity of these strains, very low Ubc9 abundance on its own is not sufficient to trigger growth defects, as we observed with the Ubc9-TO strain treated with doxycycline in which Ubc9 levels are nearly undetectable (Fig. 4). In any case, these data demonstrate that even a mild heat shock triggers elevated sumoylation in yeast, and suggest that active sumoylation, from functional levels of Ubc9, is needed for the increase in SUMO conjugation levels.

**Figure 5.**
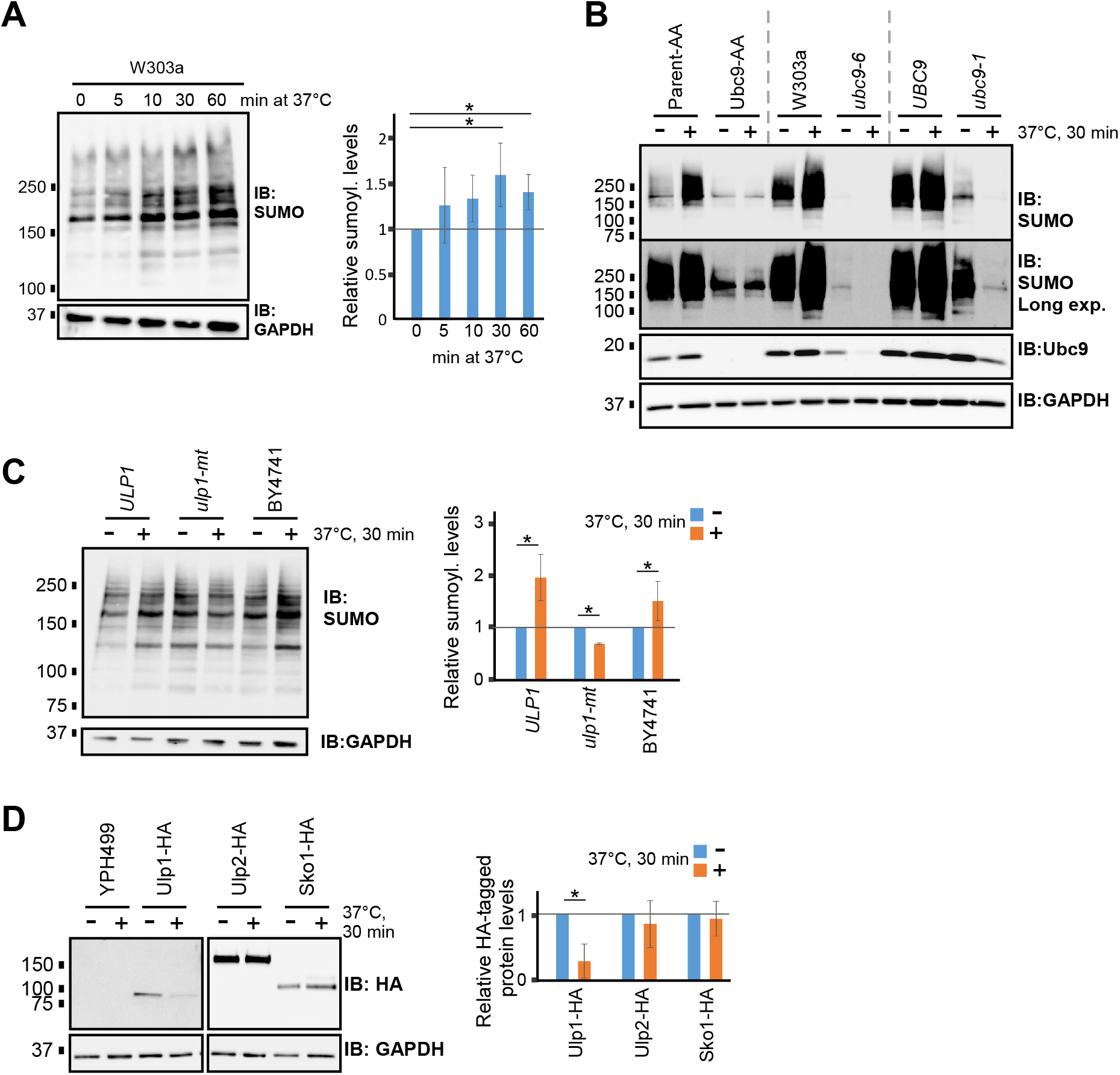
Mild heat shock elevates global sumoylation levels and reduces Ulp1 protein abundance. **(A)** Lysates were prepared from W303a yeast cultures that were heat shocked at 37°C for the indicated durations, followed by SUMO and GAPDH immunoblots. Quantification of sumoylation levels of three independent analyses was performed, with average sumoylation levels graphed and standard deviations shown. Student’s *t*-test analysis indicated that the 30- and 60-minute time-point samples are significantly elevated compared to the 0-min control (*p* < 0.05). **(B)** Cultures of the Ubc9-AA, *ubc9-6*, and *ubc9-1* strains, and their respective parental strains, were heat-shocked at 37°C for 30 min (+) or untreated (-). Lysates were prepared and analyzed by SUMO, Ubc9, and GAPDH immunoblots. **(C)** Cultures of the *ulp1-mt* strain, its isogenic parental strain (*ULP1*), and the wild-type lab strain BY4741 were heat-shocked or untreated, then analyzed by SUMO and GAPDH immunoblots. Quantification is shown at right with average relative sumoylation levels graphed with standard deviations. Student’s *t*-test analysis indicates significantly elevated sumoylation levels in the wild-type strains after heat shock, and a significant but modest reduction in the *ulp1-mt* strain. **(D)** Cultures of strains expressing HA-tagged versions of Ulp1 or Ulp2, or their parental untagged strain, YPH499, were heat-shocked or untreated. A strain expressing an HA-tagged version of an unrelated protein, Sko1, was used as a control. Lysates were analyzed by HA and GAPDH immunoblots. Quantification of HA-tagged protein levels is shown at right. Student’s *t*-tests support that heat shock significantly reduces Ulp1-HA levels but does not affect Ulp2-HA or Sko1-HA levels.

Ethanol stress triggers a marked increase in SUMO conjugation levels through nucleolar sequestration of the primary yeast SUMO protease, Ulp1 (25). To explore whether reduced desumoylation may also contribute to the increase in sumoylation levels during heat shock, we examined whether impairment of Ulp1 activity affects the heat shock-mediated increase in SUMO conjugation. Cultures of the *ulp1-mt* strain, which harbours a point mutation that reduces the function of Ulp1 (I615N; (40)), its isogenic parent strain (*ULP1*), and the wild-type lab strain BY4741 were heat shocked, then lysates were prepared and analyzed by SUMO immunoblot (Fig. 5C). As expected, the *ulp1-mt* strain showed higher sumoylation levels than its parent at the normal temperature. Whereas both wild-type strains showed the expected increase in sumoylation levels after heat shock, the *ulp1-mt* strain instead showed a modest decrease in SUMO conjugation levels. The absence of increased sumoylation in *ulp1-mt* at 37°C might reflect the already-elevated constitutive levels of SUMO conjugation in that strain. Our data, therefore, suggest that functional levels of both Ulp1 and Ubc9 are needed for the heat shock-triggered increase in SUMO conjugation levels.

Protein levels and activities of human SUMO proteases (SENPs) were previously found to be reduced by heat shock (32). We therefore examined whether Ulp1 protein levels are also affected by elevated temperature. Cultures of strains expressing C-terminally 6xHA tagged Ulp1, its untagged parental strain (YPH499), and control strains expressing 6xHA-tagged Ulp2 or the unrelated transcription factor Sko1, were grown and heat shocked at 37°C for 30 min. Lysates were prepared and examined by HA immunoblot which showed that the elevated temperature did not affect Ulp2-HA or Sko1-HA abundance but, remarkably, it caused a marked reduction in Ulp1-HA levels (Fig. 5D). Together, these data indicate that increased temperature causes elevated SUMO conjugation through active sumoylation, which requires a threshold level of functional Ubc9, and by reduced desumoylation through reduced abundance of Ulp1. If Ulp1 function is already impaired, as in the *ulp1-mt* strain, the increase in sumoylation during heat shock is not observed.

### Yeast with constitutively low sumoylation levels display altered transcription and gene expression patterns

Proteins involved in transcription, and more generally in gene expression, are amongst the most frequently identified SUMO conjugates across species (1–4, 8). To explore whether constitutively reduced levels of sumoylation affect transcription and gene expression, we carried out RNA polymerase II (RNAPII) ChIP-seq and RNA-seq analyses in the *ubc9-6* strain. Since promoter-proximal pausing is rare in yeast, the level of RNAPII occupancy on gene bodies can be a reasonable measure of transcriptional activity (41). We analyzed data from our previously reported RNAPII ChIP-seq study in which independent duplicate experiments were performed in the *ubc9-6* and W303a parental strains at the normal growth temperature, and after a 12-min heat shock at 37°C (Fig. S2A; Table S4; (16)). Comparison of RNAPII occupancy levels across all protein-coding genes revealed that, at the normal growth temperature, 736 genes showed significantly increased RNAPII occupancy whereas 846 genes had reduced RNAPII occupancy in *ubc9-6* compared to its parent (Fig. 6A; Table S4). Sample gene alignments and ChIP-qPCR validations of five genes are shown in Fig. S2B and C, respectively. Gene ontology (GO) term analysis showed that some biological processes are enriched in the subsets of genes that have elevated or reduced RNAPII occupancy in the *ubc9-6* strain (Fig. 6B; Table S5). For example, as we further discuss below, many genes involved in protein synthesis, including most ribosomal protein genes (RPGs), show significantly elevated RNAPII occupancy when sumoylation levels are low (Fig. 6B; Fig. S2B; Table S4; see below).

**Figure 6.**
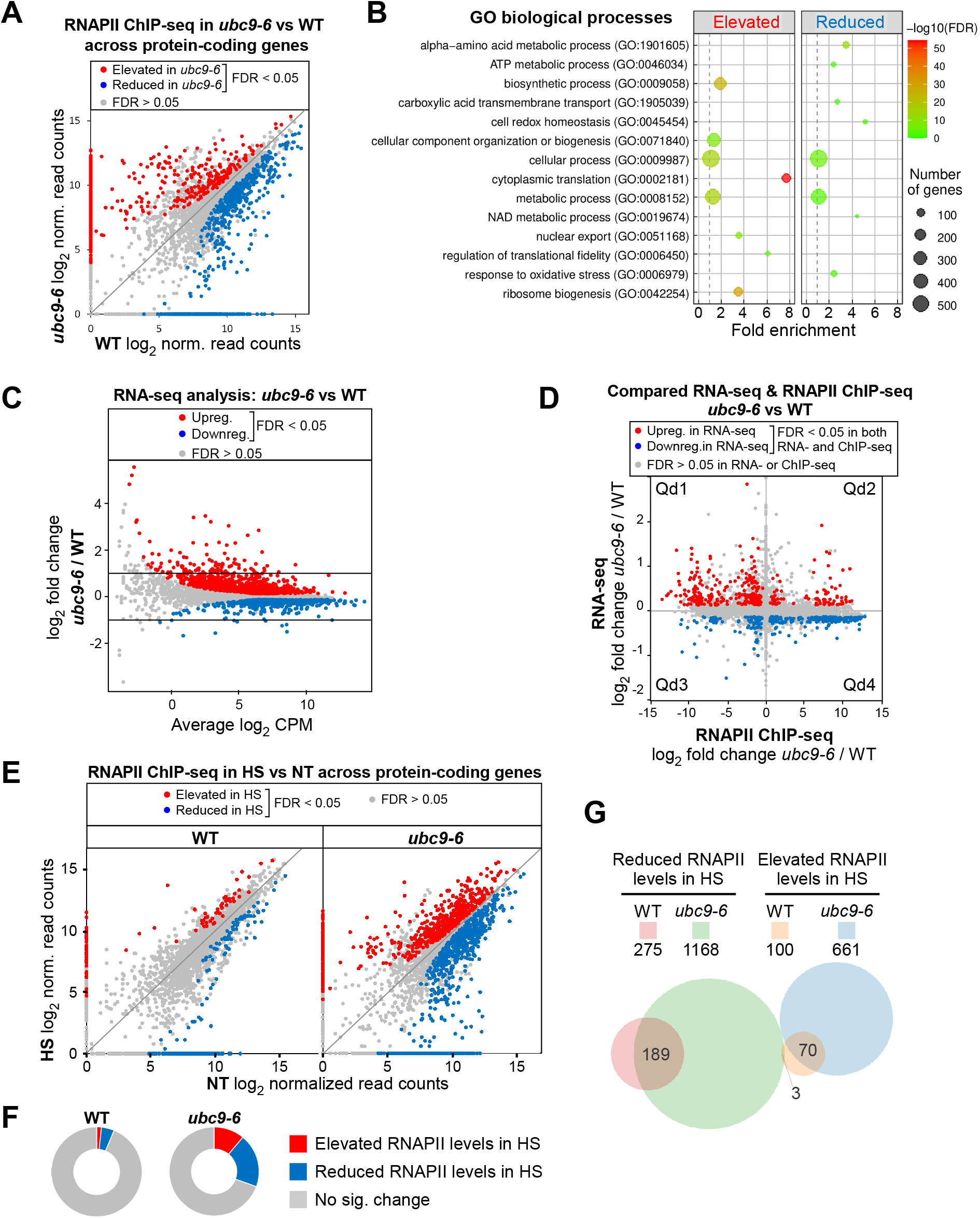
Constitutively reduced sumoylation alters gene expression patterns and expands the heat shock-induced genomic redistribution of RNAPII. **(A)** Scatterplot comparing RNAPII levels at all protein-coding genes in WT and *ubc9-6* strains, as determined by our previously reported ChIP-seq (16) (see Table S4). **(B)** GO term analysis was performed on genes showing elevated or reduced RNAPII levels across their ORFs in *ubc9-6* (see Experimental Procedures and Table S5 for details). **(C)** Comparison of global steady-state mRNA levels in *ubc9-6* and WT strains as determined by RNA-seq (see Tables S3 and S6). **(D)** Scatterplot comparing *ubc9-6*-mediated changes in mRNA abundance and RNAPII occupancy for all protein-coding genes based on the RNA-seq and RNAPII ChIP-seq analyses. Genes showing significantly altered levels of both RNAPII occupancy and mRNA abundance are shown in red or blue, as indicated (see Table S7). The four quadrants (Qd1 to Qd4) refer to subsets of genes that show correlated or anti-correlated effects in the RNA-seq and RNAPII ChIP-seq studies (see Table S7). **(E)** Scatterplots comparing heat shock-induced changes to RNAPII occupancy levels across protein-coding genes in WT and *ubc9-6* strains. RNAPII ChIP-seq was performed in both strains in untreated (NT) or heat-shocked cultures (HS; 37°C for 12 min) as previously described (16). Genes showing significantly elevated or reduced RNAPII occupancy in the heat shock samples are shown in red or blue, respectively (see Table S4). **(F)** Doughnut plots showing the proportion of protein-coding genes that display altered RNAPII levels after heat shock in WT and *ubc9-6* strains. **(G)** Venn diagram comparing the number of genes showing significantly reduced or elevated RNAPII levels after heat shock in the WT and *ubc9-6* strains.

To determine whether changes to genome-wide transcription in the *ubc9-6* strain result in corresponding altered steady-state mRNA levels, independent duplicate RNA-seq analyses were performed (Fig. S3A; Tables S3, S6). Differential expression analysis revealed that many genes showed a small, significant difference in mRNA levels in *ubc9-6* and wild-type yeast, with 700 genes upregulated and 517 genes downregulated in *ubc9-6* (Fig. 6C; Table S6). GO term analysis identified certain biological processes that are enriched among these groups (Fig. S3B). We compared our data with a previously reported RNA-seq analysis, which was performed with the *ubc9-1* strain and its parent grown in rich medium (YPD; whereas our experiments used synthetic medium, SC; (42)). As shown in Fig. S3C, even though different mutant strains and growth conditions were used, there is substantial overlap in the datasets, supporting the idea that expression of many genes is controlled by cellular sumoylation levels (Table S6). Sample gene alignments from our RNA-seq analysis and RT-qPCR validation of a subset of six genes are shown in Figs. S3D and E, respectively.

Although approximately one-fifth of all protein-coding genes showed altered expression in *ubc9-6* in our RNA-seq analysis, the majority displayed only a minor change (<2-fold; Fig. 6C). This suggests that the many changes to transcription that occur when sumoylation levels are constitutively low do not necessarily translate to corresponding changes in steady-state mRNA levels. To explore this more directly, we compared our RNAPII ChIP-seq and RNA-seq data, examining genes that are significantly altered in both datasets (referred to as “sumoylation-regulated genes”; Table S7). As shown in Fig. 6D, no clear positive correlation was observed, suggesting that sumoylation can regulate transcription and steady-state mRNA levels independently of each other. We examined the genes that lie within each of the four quadrants shown in Fig. 6D, representing subsets of sumoylation-regulated genes that are correlated or anti-correlated in the ChIP- and RNA-seq datasets, and GO term analysis identified biological processes enriched in each subset except Qd1 (Fig. S4A). Intriguingly, most sumoylation-regulated genes showed anti-correlated effects in the two studies (i.e., genes with elevated RNAPII occupancy showed reduced mRNA, and vice versa). Most notably, genes expressing translation-related proteins, including most RPGs, lie in Qd4, with increased RNAPII occupancy but modestly reduced mRNA levels, in *ubc9-6* (Fig. S4A, B). This is consistent with a previous study that showed reduced expression of RPGs in the *ubc9-1* strain (42), and suggests that a mechanism might be in place for compensating abnormally increased transcription of RPGs by decreasing stability of their mRNAs to maintain homeostasis of ribosomal protein expression. Indeed, despite elevated transcription and slightly reduced mRNA abundance, we found that levels of proteins expressed by two RPGs are unaffected in the *ubc9-6* strain (Fig. S4C).

### The heat shock-induced genomic redistribution of RNAPII is greatly expanded when sumoylation levels are low

Heat shock in yeast is accompanied by rapid changes to gene expression patterns (43–46). Considering that increased temperature also triggers elevated sumoylation, we examined whether constitutively low sumoylation levels affect the transcriptional response to heat shock. Occupancy levels of RNAPII across protein-coding gene bodies were compared in wild type and *ubc9-6* strains after a 12-min heat shock at 37°C. In the wild-type strain, a small number of genes (~6%) showed significantly increased or decreased RNAPII occupancy after the heat shock (Fig. 6E, F; Table S4). This is less than the reported ~10% - 15% of genes that show altered mRNA levels after heat shock (46), which might reflect that the expression of many heat shock-regulated mRNAs can occur post-transcriptionally (e.g., by regulating their stability), or that our differential binding analysis comparing the untreated and heat shocked RNAPII ChIP-seq datasets was highly stringent. Strikingly, however, under the same growth and analysis conditions as for the wild-type strain, the *ubc9-6* strain displayed a far more dramatic change to RNAPII occupancy levels across gene bodies (Fig. 6E, F; Fig. S4D; Table S4). Indeed, ~30% of all protein coding genes showed elevated or decreased RNAPII levels after heat shock in *ubc9-6*, which includes most of genes that show heat shock-altered RNAPII levels in the wild-type strain plus a large number of additional genes (Fig. 6G). This analysis strongly suggests that normal levels of Ubc9 activity and/or sumoylation are important for restraining the redistribution of RNAPII that occurs in response to elevated temperatures.

## DISCUSSION

A variety of environmental stressors have been shown to trigger rapid increases in SUMO conjugation levels in yeast, plants, and mammalian systems, leading to the concept of the SSR (27, 47, 48). In yeast, these include exposure to oxidative, osmotic, and ethanol stresses, high-temperature heat shock, nitrogen starvation, and DNA damaging agents (25, 26, 28, 33). We have now identified non-stress conditions that result in dramatically modulated sumoylation levels. Specifically, we found that diploid yeast harbour less sumoylation than their haploid counterparts, and that nutrient availability, or the enhancement in growth that it enables, has a positive effect on levels of SUMO conjugation. This was apparent by comparing SUMO conjugation levels in yeast grown in rich versus synthetic medium, and by monitoring sumoylation during growth in liquid culture. Sumoylation levels were greatest during exponential growth, but dropped dramatically with the diauxic shift, which occurs as glucose becomes depleted. Many proteins that become differentially sumoylated during the SSR have been identified, leading to speculation that certain processes, in particular transcription by RNAPII or RNAPIII, are influenced by altered sumoylation levels during stress (26, 28, 33). However, Lewicki et al. found that impairing transcription prior to exposing yeast to osmotic stress prevented a surge in SUMO conjugation levels, leading the investigators to speculate that the SSR is a consequence of altered transcription programs that occur during stress exposure and are not directly part of the stress response itself (26). Likewise, the elevated sumoylation that we detected in yeast during nutrient-rich and exponential growth conditions may simply be a consequence of increased rates of certain physiological processes that involve sumoylated proteins. At the same time, elevated sumoylation itself might facilitate these processes and/or high rates of growth.

Arguing against a role for high levels of sumoylation in facilitating growth, however, is our finding that yeast with dramatically reduced sumoylation levels show no apparent growth defect. Remarkably, on both solid media and in liquid culture, strains with low Ubc9 activity, either through mutation (*ubc9-1*) or reduced Ubc9 expression (Ubc9-AA, Ubc9-TO in presence of doxycycline), have normal growth rates, even when sumoylation levels are ~75% lower than in wild-type counterparts. This strongly implies that most SUMO conjugation events are dispensable under normal growth conditions, even during exponential growth where sumoylation levels normally rise. Consistent with the finding that multiple components of the sumoylation pathway are essential, including Ubc9 and Ulp1, we found that a low threshold level of sumoylation, of greater than ~3% of wild-type levels, is needed for normal growth, however (24, 49). Indeed, the *ubc9-6* strain, which shows a more dramatic reduction in sumoylation than the *ubc9-1* or untreated Ubc9-AA strains, has a modest growth defect, and the Ubc9-AA strain treated with rapamycin, which nearly abolishes sumoylation, cannot grow. This might reflect that there are essential roles for the modification in some processes, such as progress thorough mitosis and the cell-cycle (24, 49, 50). The bulk of sumoylation, however, is not needed and might be a consequence of physiological processes that involve sumoylation but do not require much sumoylation. Alternatively, as we propose below, constitutive Ubc9 activity might be required pre-emptively for protecting yeast from some, but not all, anticipated stress conditions.

Our examination of the effects of reduced sumoylation on transcription showed that about a quarter of all protein-coding genes showed significantly elevated or reduced RNAPII occupancy levels in the *ubc9-6* strain compared to its wild-type parent. This indicates that reduced sumoylation has a widespread effect on transcription by RNAPII and implies that normal levels of sumoylation are needed to maintain global transcriptional patterns. Transcriptome analyses in human cells in which SUMO1 or Ubc9 expression levels were reduced by RNA interference showed that steady-state mRNA levels of a variety of gene classes were upregulated or downregulated when sumoylation levels dropped (13, 14). Our RNA-seq analysis in the *ubc9-6* strain, however, showed only modest changes to the mRNA levels of a number of genes compared to wild-type, which is consistent with a previous study using the *ubc9-1* mutant which found that the transcriptome was barely affected by the mutation (42). In yeast with constitutively low sumoylation, then, it is possible that an adaptive mechanism is in place to buffer changes in transcription, perhaps by controlling RNA degradation rates correspondingly, such that the transcriptome is mostly unaffected. Supporting this, we find that most of the small but significant changes to mRNA levels in *ubc9-6* anti-correlate with changes to RNAPII occupancy on the corresponding genes, as though cells are working to elevate mRNA stability for genes with abnormally reduced transcription levels, and reduce stability for mRNAs of genes with abnormally elevated transcription. Indeed, systems for buffering changes in mRNA synthesis, due to perturbation of transcription, with reciprocal changes to mRNA degradation have been described (51, 52).

The rise in SUMO conjugation that occurs as part of the SSR after exposure to a variety of stress conditions suggests that sumoylation plays a protective role or facilitates the stress response (17, 27). It is surprising then, that we observed no growth defect in yeast with low Ubc9 activity and reduced sumoylation levels that were exposed to oxidative, osmotic, or ethanol stress. Our observations align with the proposal that the SSR is primarily a consequence of altered transcription genome-wide, which involves a wave of transcription-mediated SUMO conjugation, and suggest that high levels of overall sumoylation do not play a protective role during these particular stresses (18, 26). In contrast, all tested strains with reduced Ubc9 activity, and consequently low sumoylation levels, showed growth defects when incubated at elevated temperatures, with most showing severe growth impairment. This is consistent with the finding that sumoylation facilitates heat tolerance in human cells and plants (31, 53, 54). For example, siRNA-mediated depletion of SUMO1 and SUMO2 levels resulted in a seven-fold reduction in survival after heat shock in human U2OS cells (31). Heat shock in human cells leads to increased polysumoylation of proteins involved in multiple processes, including transcription, and it triggers altered sumoylation of promoter- and enhancer-bound proteins, specifically (17, 31). These findings suggest that the elevated sumoylation that occurs with high temperature can function, at least in part, to adjust transcriptional programs as needed for surviving heat stress.

We explored this by examining how reduced constitutive sumoylation levels affect the transcriptional response to heat shock. In yeast, 10% - 15% of genes show upregulated or downregulated mRNA levels in response to heat shock (43, 44, 46). Although heat shock can change steady-state mRNA levels through regulation of mRNA stabilities, much of the heat shock-altered transcriptome is likely due to changes at the transcriptional level (55). Indeed, significant reorganization of multiple transcription-related factors on chromatin has been observed in quick response to elevated temperature (46). In our analysis, ~6% of protein-coding genes showed altered RNAPII occupancy levels in wild-type cells after heat shock, but this number increased dramatically to ~30% in the *ubc9-6* strain. This strongly implicates Ubc9 and sumoylation in controlling or restraining the transcriptional response to elevated temperatures. More specifically, we believe that a rise in SUMO conjugation levels, as opposed to merely a threshold constitutive level of sumoylation, is needed to endure high temperatures, at least partly by regulating transcription. Supporting this, we observed that active sumoylation (i.e., functional levels of Ubc9) is required, but also that Ulp1 protein levels drop significantly after heat shock, implying that reduced desumoylation plays a major, evolutionarily conserved role in elevating sumoylation levels when temperature rises (32). Taken together, our results demonstrate that SUMO conjugations, although plentiful, are largely dispensable for normal growth in many stress and non-stress conditions. However, we find that active sumoylation and desumoylation systems are critical for enduring elevated temperatures where they function, at least in part, to temper the transcriptional response to heat shock. Further study in this area will be invaluable for uncovering the molecular mechanisms by which sumoylation facilitates heat tolerance in eukaryotes.

## EXPERIMENTAL PROCEDURES

### Yeast media and growth

Yeast strains used in this study are listed in Table S1. Previously unreported strains were generated by transformation with PCR-amplified DNA fragments that were incorporated at specific genomic loci through homologous recombination (56). Unless otherwise noted, yeast were grown at 30°C in synthetic complete (SC) medium. Overnight cultures were diluted to an absorbance (595 nm) of ~0.2 in a volume of 10 to 50 mL, then grown to exponential phase (absorbance < 1.0) unless otherwise noted, treated as indicated below if appropriate, then harvested by centrifugation at 3000 *g* for 5 min. Where appropriate, the following were added to media (final concentrations indicated): 0.6 M KCl for 5 min for liquid cultures, 1 M KCl on solid media; 1M NaCl; 100 mM H2O2 for 5 min for liquid cultures, 1 mM H2O2 for solid media; 10% ethanol for 60 min for liquid cultures, 7% ethanol for solid media; 1 μg/mL rapamycin for 30 min, or as indicated; doxycycline, 10 μg/mL, or as indicated, for duration of culture growth. Where rapamycin was added, the control or “untreated” sample was supplemented with the same volume of its solvent, DMSO.

### Protein lysates and extracts

Protein samples were generated under non-denaturing conditions (“lysates”) or through precipitation with trichloroacetic acid (TCA; “protein extracts”). For lysate preparation, pellets were washed with ice-chilled IP buffer (50 mM Tris-HCl, pH8; 150 mM NaCl; 0.1% Nonidet P-40 (NP40); supplemented with a protease inhibitor cocktail, 1 mM phenylmethylsulphonyl fluoride (PMSF), and 2.5 mg/mL *N*-ethylmaleimide (NEM)) at a volume equal to the culture volume, then resuspended in 500 μL of IP buffer. Acid-washed glass beads (0.25 g) were then added, and samples were vortexed at 4°C for 30 min, with a 5-min ice incubation after the first 15 min. Lysates were transferred to fresh microfuge tubes and clarified by two rounds of 5-min 4°C centrifugation at maximum microfuge speed. Prior to analysis by immunoblot, an equal volume of 2X sample buffer (140 mM Tris-HCl, pH 6.8; 4% SDS; 20% glycerol; 0.02% bromophenol blue; supplemented with 10% 2-mercaptoethanol prior to use) was added, and samples were boiled for 4 min.

For protein extract preparation through TCA precipitation, volumes equal to 2.5 absorbance units (595 nm) of the exponentially growing yeast cultures, treated if necessary, were centrifuged at 3000 *g*, then resuspended in 1 mL of ice-cold water. 150 μL of freshly prepared extraction buffer (1.85 NaOH; 7.5% 2-mercaptoethanol) was added, and the mixed samples were incubated on ice for 10 min. Then, 150 μL of cold 50% TCA was added, samples were mixed, then incubated again on ice for 10 min before they were centrifuged at 4°C at maximum microfuge speed. Supernatants were removed and the protein pellets were resuspended in 100 μL of 2X sample buffer, with 5 to 10 μL of Tris-HCl, pH 8 added if the sample was yellow in colour, to raise the pH and restore the blue colour. Samples were boiled for 4 min, centrifuged for 3 min at maximum microfuge speed to separate debris, then used for immunoblot analysis.

### Immunoblots and quantifications

Protein samples were analyzed by standard polyacrylamide gel electrophoresis (PAGE) techniques. For preferential detection of high-molecular weight SUMO conjugates, PAGE was performed with lysates followed by wet transfer (100 V for 1 hr) in a buffer consisting of 0.1% SDS, 10% methanol, 50 mM Tris-HCl, and 380 mM glycine. For preferential detection of free SUMO and lower molecular-weight SUMO conjugates, TCA extracts were analyzed by PAGE, followed by semi-dry transfer using a Power Blotter system (ThermoFisher). Chemiluminescence-based imaging was performed using the MicroChemi imager (DNR). For quantification of signals, TIFF-format images were analyzed using ImageJ software (version 1.52a; NIH). Specifically for determining sumoylation levels, all SUMO signals (above the Ubc9-SUMO band, if it was present) were considered SUMO conjugates, and normalization was made to the corresponding signal from GAPDH immunoblots. Antibodies used for immunoblot analyses were: 1:500 – 1:1000 SUMO/Smt3 (y-84; Santa Cruz, sc-28649); 1:3000 GAPDH (Sigma, G9545); 1:500 Ubc9 (Santa Cruz, sc-6721); 1:3000 histone H3 (Abcam, ab1791); 1:1000 Myc epitope tag (Sigma, 05-724); 1:1000 Rpb3 (Abcam, ab202893); 1:5000 HA (12CA5; Sigma, 11583816001).

### Spot assays and liquid growth assays

Yeast strain fitness was examined by spot assays, as previously described (57), or liquid growth analysis. To generate liquid growth curves, growing cultures were diluted to an absorbance of 0.2 – 0.5 in appropriate medium and treated if needed. Each sample was transferred, in triplicate, to wells in a 96-well microtiter dish, and inserted into an accuSkan absorbance microplate reader (Fisherbrand), which shook and incubated samples at the appropriate temperature while taking absorbance measurements (595 nm) at 15-min intervals over a course of 20 hours. Background absorbance readings were made from wells containing uninoculated medium, and the background-subtracted average absorbance readings, and standard deviations, are listed in Table S8. Curves shown in figures were generated using average values without standard deviations indicated since they are minimal and would otherwise obscure curves on the diagrams. Because the light path distance in the accuSkan is less than 1 cm, absorbance readings are indicated as arbitrary units (a.u.) and are not directly comparable to values obtained through a standard spectrophotometer.

### RNA-seq and RNAPII ChIP-seq analysis

RNA preparation and ChIP were performed as previously described (16). Details of the *ubc9-6* RNA-seq experiment are presented in Table S3. RNAPII ChIP-seq analysis was previously reported (16), with the heat shock-treated samples (37°C for 12 min) analysed in the same manner. Bioinformatics analysis of the previously published *ubc9-1* RNA-seq dataset (42) was also performed as indicated in Table S3. Read counts and differential expression analysis results for the RNA-seq analyses are in Table S6, and RNAPII ChIP-seq analysis results are in Table S4. GO term analysis and visualization were performed as previously described (58) with data shown in Table S5. Validation qRT-PCRs were performed as previously described using primers indicated in Table S2 (16).

## DATA AVAILABILITY

Data for the RNA-seq analysis with WT and *ubc9-6* strains were deposited in the Gene Expression Omnibus (GEO) repository with accession number GSE167427. The GEO accession number for the previously reported RNAPII ChIP-seq with the same strains is GSE167424.

## ACKNOWLEDGEMENTS

This work was supported by grants from the Canadian Institutes of Health Research (CIHR; grant numbers MOP-142282 and PJT-178112) and the Natural Sciences and Engineering Research Council of Canada (NSERC; RGPIN-04208-2014) to E.R.

## SUPPORTING INFORMATION FIGURE LEGENDS

### Supplementary Figures (PDF format)

**Figure S1. Detection of Ubc9-SUMO and characterization of the Ubc9-AA strain. (A)** The Ubc9-SUMO species persists in highly reducing conditions implying that it does not derive from a thioester linkage but represents covalently sumoylated Ubc9. Lysates from W303a and 8HIS-Smt3 strains were prepared in sample buffer containing the indicated concentrations of β-mercaptoethanol (β-me), then analyzed by PAGE and SUMO and GAPDH immunoblots. **(B)** Although it harbours constitutively lower SUMO conjugation levels, the Ubc9-AA strain shows increased sumoylation levels in ethanol, oxidative, and osmotic stress conditions, as previously shown for WT yeast (see text). The Ubc9-AA strain was grown in the indicated conditions (see Experimental Procedures for details), then lysates were prepared and analyzed by SUMO and GAPDH immunoblot.

**Figure S2. Analysis and validation of RNAPII ChIP-seq in the *ubc9-6* strain. (A)** Cluster analysis showing similarity of two independent replicates of our previously reported RNAPII ChIP-seq analysis in WT and *ubc9-6* strains (see Table S4 for data; (16)). **(B)** Sample alignments of independent duplicate RNAPII ChIP-seq analyses using the Integrative Genomics Viewer (IGV) genomic alignment tool. Reads are not normalized, and values shown correspond to maximum data range (read numbers) for the view shown. Arrows indicate position and direction of ORFs for selected genes. **(C)** RNAPII ChIP-qPCR validation of ChIP-seq data. qPCR was performed using primers specific to the indicated genes (see Table S2 for PCR primer sequences) and templates obtained from at least three independent RNAPII ChIP experiments. Average folds over background are graphed with standard deviations shown. Student’s *t*-test was performed to assess the statistical significance of changes to levels in WT versus *ubc9-6* strains.

**Figure S3. Analysis and validation of RNA-seq in the *ubc9-6* strain. (A)** Cluster analysis showing similarity of two independent replicates of RNA-seq analysis in WT and *ubc9-6* strains (see Tables S3 and S6). **(B)** GO term analysis was performed on differentially expressed up- or downregulated genes in *ubc9-6* (see Experimental Procedures and Table S5 for details). **(C)** Comparison of *ubc9-6*- and *ubc9-1-specific* changes to mRNA levels. Fold changes of global mRNA levels were compared in our RNA-seq analysis of *ubc9-6* (grown in SC medium) and in a published RNA-seq analysis of *ubc9-1* (grown in YPD medium; European Nucleotide Archive accession PRJEB7579; (42)). Comparison is shown as scatterplot with Pearson correlation coefficient, *r*, indicated. Middle, doughnut charts showing the fraction of genes up- or downregulated in *ubc9-6* that were regulated or not regulated in the *ubc9-1* analysis. At right, Venn diagram comparing number of genes up- or downregulated in the *ubc9-6* and *ubc9-1* analyses. **(D)** Sample alignments of independent duplicate RNA-seq analyses and the RNAPII ChIP-seq analysis using the Integrative Genomics Viewer (IGV) genomic alignment tool. Reads are not normalized. Values shown correspond to maximum data range (read numbers) for the view shown and arrows indicate position and direction of ORFs for selected genes. **(E)** RT-qPCR analysis of indicated genes from RNA isolated from *ubc9-6* or WT strains (compare with *D*). Average mRNA levels, relative to WT levels, are shown. Error bars represent standard deviation of three replicates and asterisks (*) indicate Student’s *t*-test *p*-values of < 0.05. Although the *t*-test does not indicate a statistically significant difference for *HSP26*, mRNA levels are notably elevated in each replicate, but to different degrees, inset. Sequences of PCR primers used are indicated in Table S2

**Figure S4. Additional RNAPII ChIP-seq, RNA-seq, and RPG analyses in the *ubc9-6* strain. (A)** Genes that showed significantly altered levels in both RNAPII occupancy and mRNA abundance in the *ubc9-6* strain (compared to WT) were categorized into four subsets based on how they correlated in the two analyses. Each subset is shown as a quadrant (Qd1 to Qd4) in Figure 6D and listed in Table S7. GO analysis was performed on genes from each quadrant as described in Experimental Procedures and in Table S5. No significant GO terms were identified for genes in Qd2. **(B)** Scatterplot showing the behaviour of all ribosomal protein genes (RPGs) in our RNA-seq and RNAPII ChIP-seq analyses. Many show significantly elevated RNAPII occupancy levels but slightly reduced mRNA abundance in the *ubc9-6* strain compared to WT. **(C)** The *ubc9-6* mutation does not affect protein levels for RPGs. Strains were generated expressing 3xHA C-terminal tagged versions of RPL5 and RPS20 from their natural loci in WT or *ubc9-6* backgrounds. Protein extracts were prepared from cultures of these strains and analyzed by HA and GAPDH immunoblots. **(D)** Sample alignments of RNAPII ChIP-seq analyses in WT and *ubc9-6* strains in untreated (NT) and heat shock (HS) conditions using the Integrative Genomics Viewer (IGV) genomic alignment tool. Reads are not normalized. Values shown correspond to maximum data range (read numbers) for the view shown and arrows indicate position and direction of ORFs for selected genes.

### Supplementary Tables (PDF format)

**Table S1.**
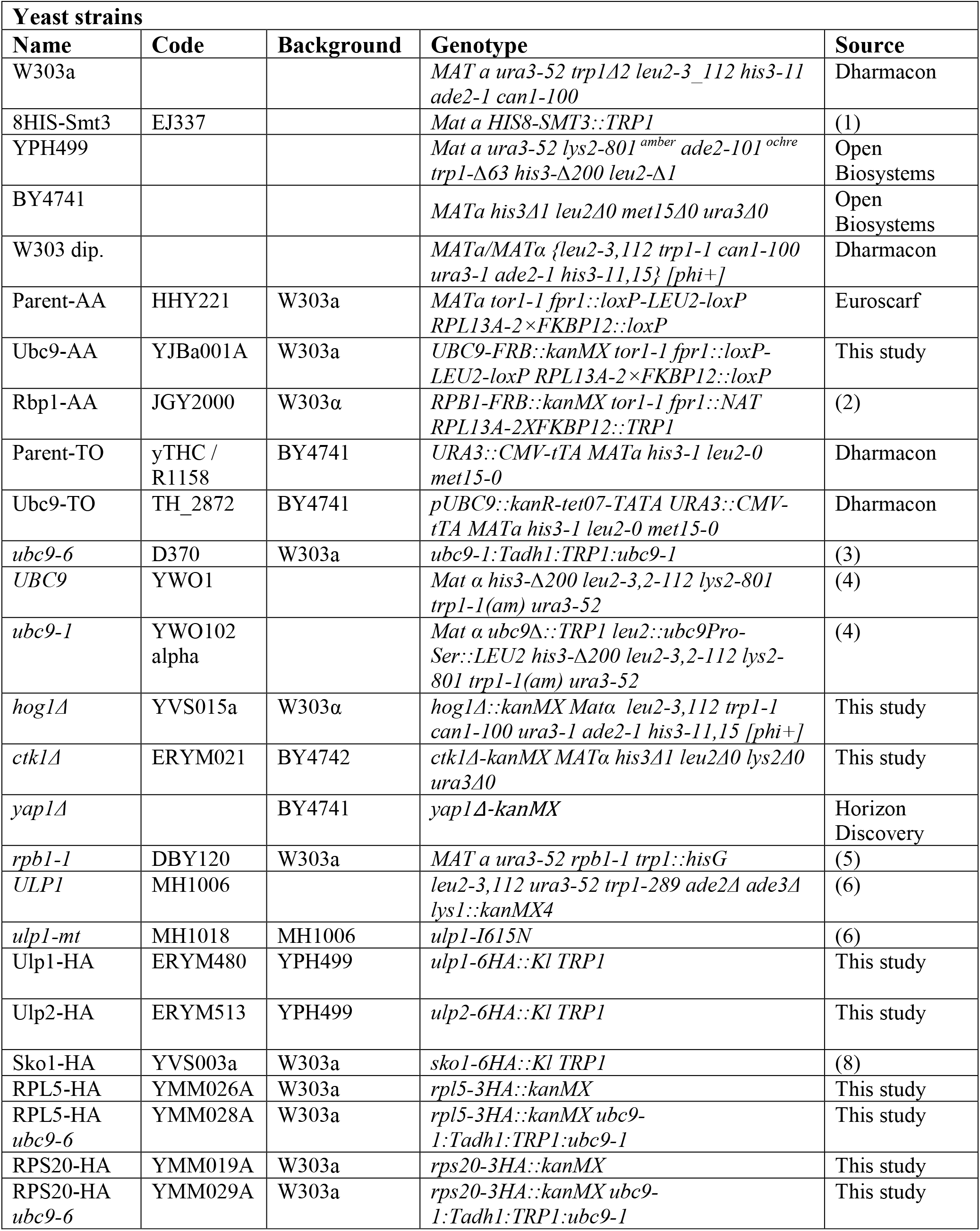
Yeast strains used in this study. **REFERENCES** 1. Wohlschlegel JA, Johnson ES, Reed SI, Yates 3rd JR. 2004. Global analysis of protein sumoylation in Saccharomyces cerevisiae. J Biol Chem 279:45662–45668. 2. Fan X, Moqtaderi Z, Jin Y, Zhang Y, Liu XS, Struhl K. 2010. Nucleosome depletion at yeast terminators is not intrinsic and can occur by a transcriptional mechanism linked to 3’-end formation. Proc Natl Acad Sci U S A 107:17945–50. 3. Baig MS, Dou Y, Bergey BG, Bahar R, Burgener JM, Moallem M, McNeil JB, Akhter A, Burke GL, Sri Theivakadadcham VS, Richard P, D’Amours D, Rosonina E. 2021. Dynamic sumoylation of promoter-bound general transcription factors facilitates transcription by RNA polymerase II. PLOS Genet 17:e1009828. 4. Seufert W, Futcher B, Jentsch S. 1995. Role of a ubiquitin-conjugating enzyme in degradation of S- and M-phase cyclins. Nature 373:78–81. 5. McNeil JB, Agah H, Bentley D. 1998. Activated transcription independent of the RNA polymerase II holoenzyme in budding yeast. Genes Dev 12:2510–2521. 6. Soustelle C, Vernis L, Freon K, Reynaud-Angelin A, Chanet R, Fabre F, Heude M. 2004. A new Saccharomyces cerevisiae strain with a mutant Smt3-deconjugating Ulp1 protein is affected in DNA replication and requires Srs2 and homologous recombination for its viability. Mol Cell Biol 24:5130– 5143.

**Table S2.**
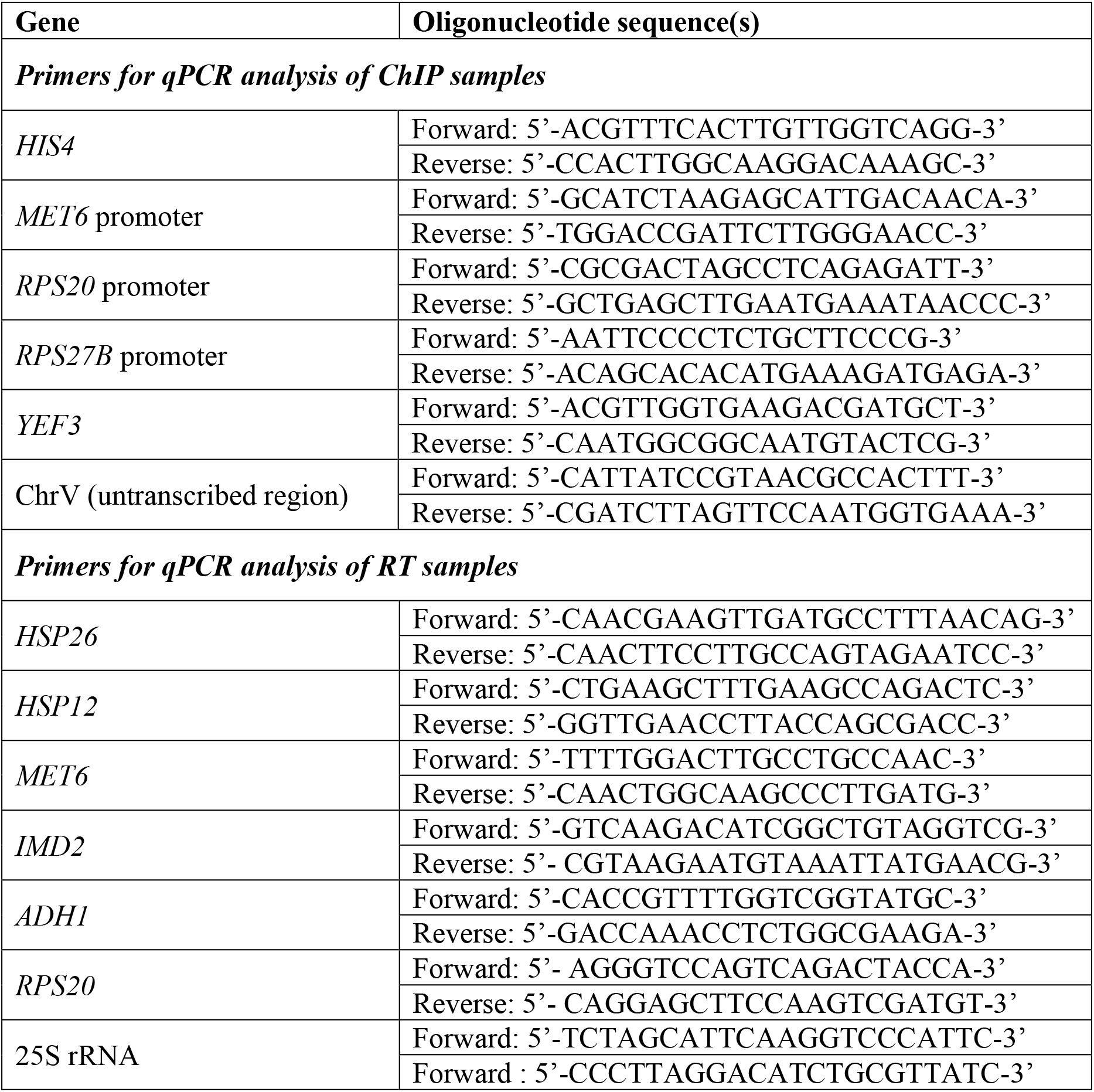
Sequences of oligonucleotides used in this study.

**Table S3.** Analysis details for RNA-seq in *ubc9-6* vs. WT.

| RNA-seq in *ubc9-6* and WT strains |  |
| --- | --- |
| Samples and conditions | Two independent replicates were prepared from cultures grown in SC medium at 30°C.<br>1 – W303a (“WT”)<br>2 – *ubc9-6* |
| Library synthesis | NEB Ultra II directional mRNA library prep (PolyA enrichment) |
| Sequencing | Illumina HiSeq 2500; Paired-end reads; 2x 126 nt; 30 million reads/sample |
| Quality control | FastQC (0.11.9) |
| Genome alignment | HISAT2 (2.1.0)<br>**Parameters:** Default options; UCSC annotation file: sacCer3.ncbiRefSeq.gtf |
| Transcript counting | Transcripts counted using featureCounts from Subread (2.0.0)<br>**Parameters:** Default options; -p (paired) |
| Differential expression analysis | edgeR (3.32.1)<br>**Parameters:** Library sizes were normalized using calcNormFactors; dispersions were estimated with estimateDisp with robust argument; likelihood ratio tests for differential expression were performed with glmFit and glmLRT. |
| Data availability | NCBI GEO database accession: GSE167427 |

### Supplementary Tables (Spreadsheet format)

Table S4 RNAPII ChIP-seq data analysis

Table S5 GO term analysis data

Table S6 RNA-seq data analysis

Table S7 RNAPII ChIP-seq and RNA-seq comparison data

Table S8 Data for graphs presented in figures

